# Dissociation of fear initiation and maintenance by breathing-driven prefrontal oscillations

**DOI:** 10.1101/468264

**Authors:** Sophie Bagur, Julie M. Lefort, Marie M. Lacroix, Gaëtan de Lavilléon, Cyril Herry, Clara Billand, Hélène Geoffroy, Karim Benchenane

## Abstract

Does the body play an active role in emotions? Since the original James/Cannon controversy this debate has mainly been fueled by introspective accounts of human experience. Here, we use the animal model to demonstrate a physiological mechanism for bodily feedback and its causal role in the stabilization of emotional states. We report that during fear-related freezing mice breathe at 4Hz and show, using probabilistic modelling, that optogenetic perturbation of this feedback specifically reduces freezing maintenance without impacting its initiation. This rhythm is transmitted by the olfactory bulb to the prefrontal cortex where it organizes neural firing and optogenetic probing of the circuit demonstrates frequency-specific tuning that maximizes prefrontal cortex responsivity at 4Hz, the breathing frequency during freezing. These results point to a brain-body-brain loop in which the initiation of emotional behavior engenders somatic changes which then feedback to the cortex to directly participate in sustaining emotional states.

---

The link between brain and body in emotion has been debated since the original James-Cannon controversy(1–3). The key question under dispute is whether somatic changes associated with emotion are merely the expression of an emotional state or if they feedback to the brain to play an active role, in particular in generating the subjective feel of a particular emotion. Changes in somatic physiology are strongly correlated with emotional state (3–5) and more importantly they show evidence of influencing various aspects of emotional cognition(6–8). However, the mixed and often subtle effects of the lack of bodily feedback on emotions, as initially highlighted by lesion studies in animal separating the viscera from the brain and then in human studies of spinal cord lesion patients, is a strong challenge to the claim that somatic modifications have an active role to play(9–12). Any such role is therefore unlikely to be in the genesis of emotion itself but instead will probably be restricted to the regulation of more subtle aspects of emotional cognition. Identifying these precise parameters is crucial to elaborate a mechanistic account of emotions. In particular, fear produces a strongly stereotyped behavioral state that is well characterized and readily triggered in human and non-human animals. We therefore address these issues using the canonical paradigm of rodent fear conditioning(13).

Although most research has focused on the cardiovascular system, the respiratory rhythm is a strong candidate to bridge the gap between brain and body during emotion. Breathing is strongly modulated during fear and other emotions(14,15) and numerous studies have shown that the respiratory rhythm can influence neuronal dynamics, mainly via feedback from the olfactory system(16), in many brain areas such as the hippocampus, amygdala and prefrontal cortex in both rodents(17–20) and humans(21). We hypothesized that breathing is therefore a marker of emotional state poised to directly impact neuronal functioning in the cortical and limbic system to regulate fear expression (FigS1). To test our hypothesis, we studied the mechanism and role of breathing feedback during fear related behavior in mice at the behavioral and electrophysiological level.

We conditioned mice using a classical auditory conditioning paradigm and in test sessions 24 hours later, they demonstrated robust freezing in response to presentation of the conditioned stimulus (CS+) but not the unconditioned stimulus (CS-) (Fig1A). Using a plethysmograph, we found that during active exploration of the environment, mice breathe irregularly at 8-12Hz as previously described(22)(Fig1B,C). During freezing, the characteristics of breathing were deeply modified (Fig1B): the frequency dropped to 4Hz (Fig1C) (23), the tidal volume increased relative to non-freezing periods (Fig1D) and the variability both of frequency and tidal volume dropped (Fig1E,F). Therefore, during freezing, breathing shifts to a slower, deeper and more regular 4Hz rhythm.

**Figure 1.**
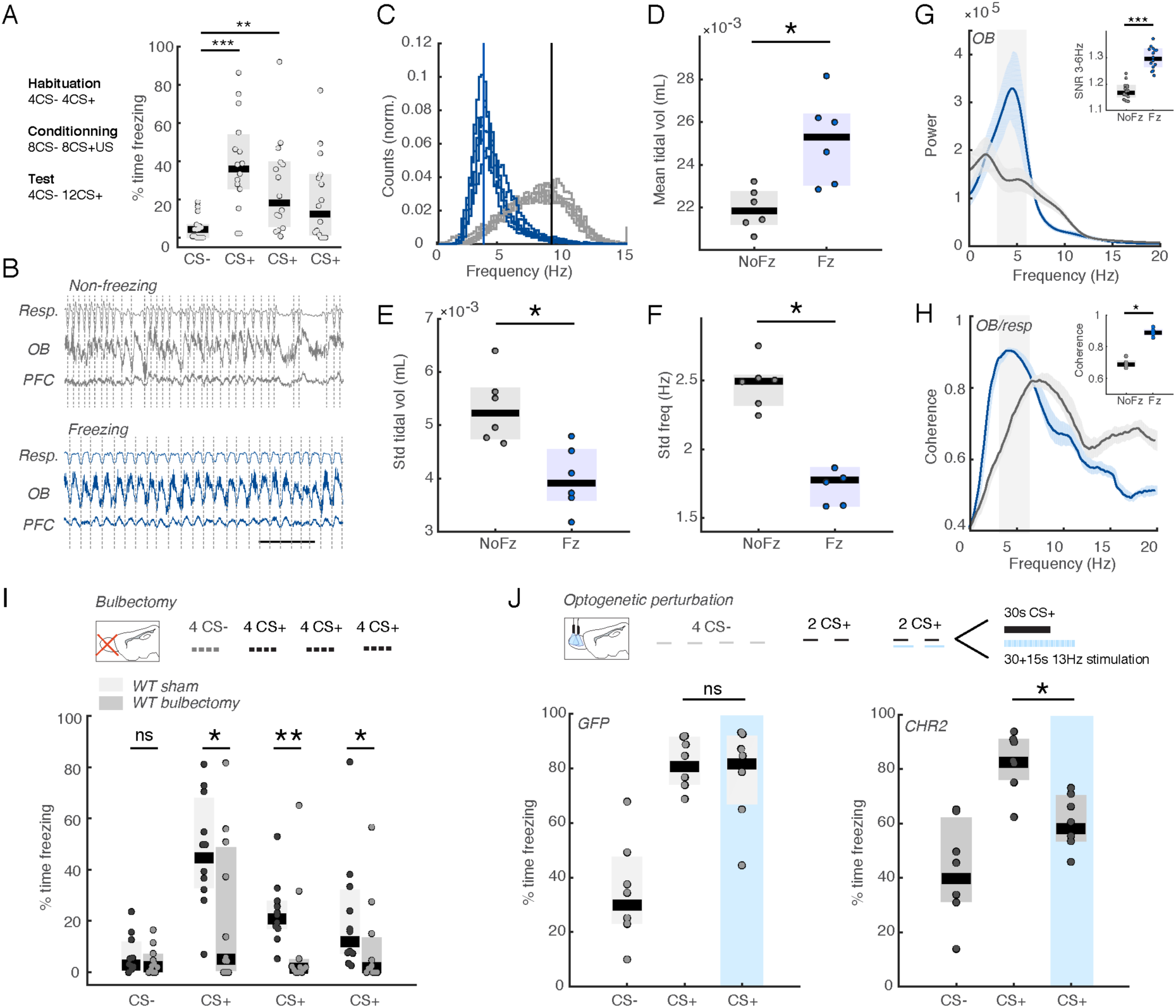
4Hz respiratory rhythm plays a crucial role in fear behaviour. A. Experimental protocol and behavioural results: median percentage of time spent freezing for all mice during the CS- and CS+, grouped as blocks of four sounds. Sound block significantly affected freezing rates (Friedman test: χ^2^(3)=19.6, p=2.01e-4). Post-hoc analysis revealed significant differences of the CS+ blocks compared to the CS-block (Wilcoxon signed rank test: Signed Rank statistic=0,14,23; p=6e-5, 0.0067, 0.0353, n=15). B. Illustrative example of breathing activity as recorded in the plethysmograph and OB and PFC LFPs during exploratory and freezing periods. Note the striking regularity and synchronicity in the 3-6Hz range in all three measures during freezing. Bar:1sec. C. Distribution of breathing frequencies recorded in the plethysmograph during freezing and non-freezing periods. (n=6) D. Mean breathing tidal volume during freezing and non-freezing periods. (Paired Wilcoxon signed rank test: Signed Rank statistic=0, p=0.0312, n=6) E.F. Variability of breathing frequency and tidal volume during freezing and non-freezing periods. (Paired Wilcoxon signed rank test: Signed Rank statistic=0,0 p=0.0312, 0.0312, n=6) G. Averaged olfactory bulb power spectra during freezing (blue) and non-freezing (gray) periods. Error bars are SEM. Inset: average signal-to-noise ratio of 3-6Hz band during freezing and non-freezing periods. The signal to noise ratio is defined as the ratio between power in the band of interest to the power in the rest of the spectrum. (Paired Wilcoxon signed rank test: Signed Rank Statistic=0, p=6.10e-4, n=15). H. Averaged coherence between breathing and the olfactory bulb during freezing (blue) and non-freezing (gray) periods. Error bars are SEM. Inset: average coherence of 3-6Hz band during freezing and non-freezing periods. (Paired Wilcoxon signed rank test: Signed Rank statistic=0, p=0.0312, n=6) I. Median freezing levels of sham and bulbectomized mice during the test session. (Two-way mixed repeated measures anova: group (p=0.041, F=4.73) x CS block (p<0.0001, F=16.28), interaction (p=0.09, F=2.18). Post-hoc Wilcoxon ranksum on each individual block: zval=0.82, 2.1, 2.6296, 2.2389; p=0.41, 0.05, 0.01, 0.02, n=11,11; Effect size for significant differences: 0.95,0.79, 0.56) J. Median freezing levels of GFP-expressing controls and ChR2-expressing mice during the test session. (Wilcoxon signed rank test: Signed Rank statistic=16, 28, p=0.813, 0.016, n=7,7; Effect size for CHR2 with/without stimulation: 2.08; Effect size CHR2 vs GFP during stimulation: 1.2)

The main feedback pathway of the respiratory rhythm is the olfactory bulb (OB) which, as expected (24), showed a prominent 4Hz oscillation during freezing that was highly coherent with respiration (Fig1G,H). If the respiratory rhythm is indeed a form of somatic feedback that regulates emotional expression as we postulated, OB ablation or perturbation will impair this feedback and should modify emotional expression. We indeed found that bilateral bulbectomy reduces freezing levels during test sessions (Fig1I). This decrease of freezing cannot be attributed to other behavioral impairments linked to bulbectomy (FigS2). Next, using optogenetics, we selectively perturbed the temporal organization of the OB activity at 4Hz by rhythmically activating interneurons (ChR2 in GAD-Cre mice) of OB glomeruli which are responsible for the low-frequency components of OB activity (25) at a competing frequency of 13HZ to scramble the OB output around the CS+ presentation (Fig1J). The stimulation reduced freezing levels in ChR2 mice but did not impact GFP controls (Fig1J). These results show that the 4Hz respiratory rhythm, transmitted via the OB, plays a crucial role in fear behaviour.

The finding that the perturbation of respiratory rhythm feedback decreases freezing leaves open the question of how this feedback impacts neural control of the emotional response. Close examination of the link between the 4Hz oscillation and behavior shows that during a freezing episode power gradually rises at freezing onset but drops off sharply at offset (Fig2A), indicating that although 4Hz power correlates with both freezing onset and offset, it is most closely associated with episode termination. Indeed, between individual animals, offset dynamics were more robust and reproducible than onset dynamics (Fig2B,C). Accordingly, 4Hz power was a better predictor of freezing vs active state around offset than around onset (FigS3). Crucially, we found that this asymmetry between freezing onset and offset cannot be explained by a difference in the behavior of the animal as measured using head acceleration (Fig2D,E).

**Figure 2.**
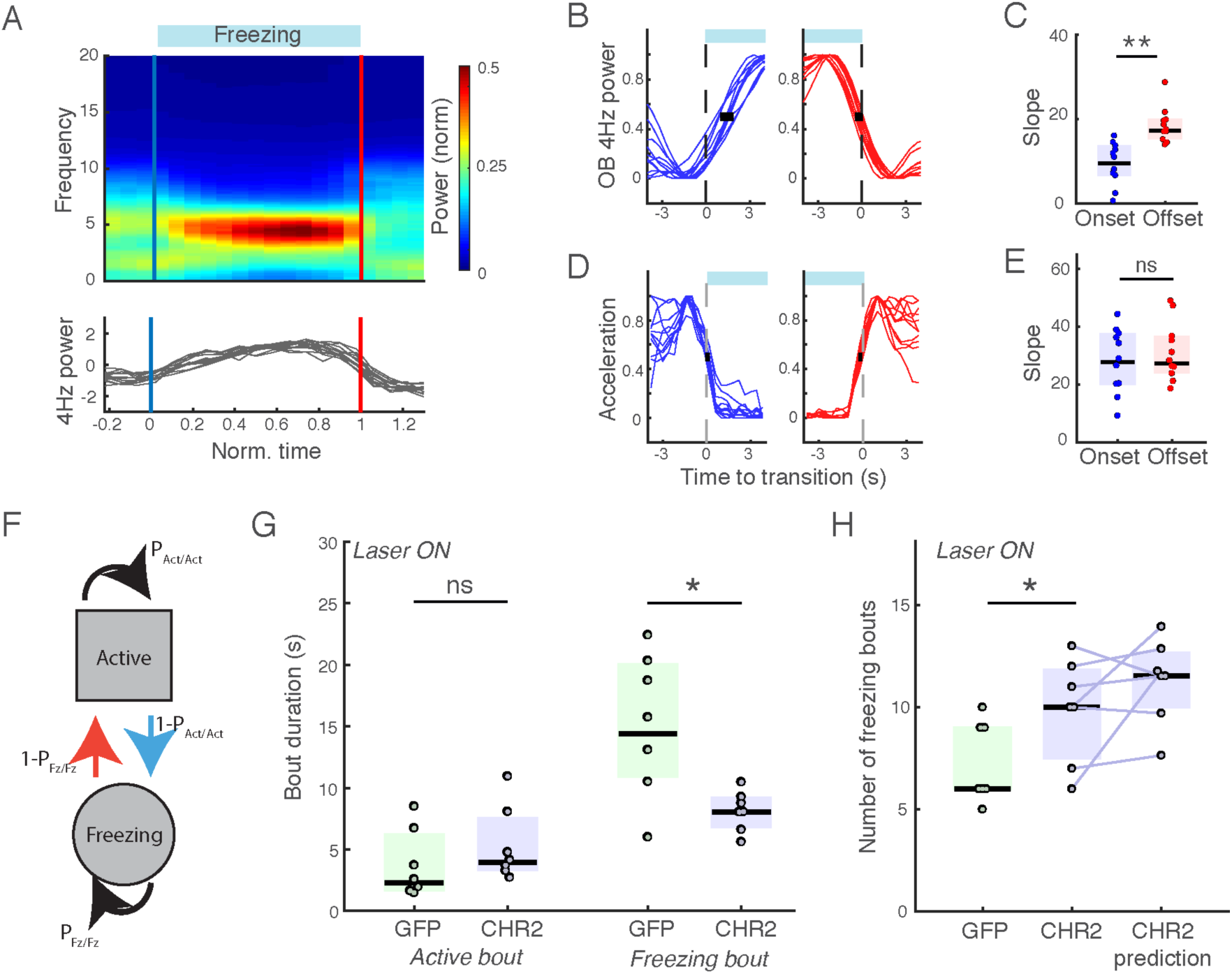
4Hz respiratory rhythm is specifically involved in the regulation of freezing maintenance. A. OB spectrogram (top) and mean power in the 4Hz band (bottom) triggered on freezing periods of normalized duration for all mice. Note the asymmetric rise and fall of 4Hz power during the freezing bout with a gradual onset and steep offset. (n=15 mice) B. D OB 4Hz power (B) and head acceleration (D) triggered on onset (left, blue) and offset (right, red) of freezing bouts. Black bar indicates the standard deviation at the midpoint. (n=11,11 mice, only mice with sufficient number of 4s bouts of freezing are retained). C. E Slope of onset and offset curves of and OB 4Hz power (C) head acceleration (E) to quantify the difference in transition speed, estimated using a sigmoid fit. (Wilcoxon ranksum: zval=-0,19, −4.31; p=0.19, 1.6e-5, n=11) F. Schematic of Markov model used to model freezing. At each time step, depending on the current state (Fz or Act), the next state is randomly selected according to fixed probabilities. G. Duration of active and freezing bouts and frequency of freezing bout initiation for GFP and CHR2 expressing mice during 13Hz laser stimulation in the test session. (Wilcoxon ranksum test: ranksum value=42, 71,: p=0.2, 0.017; n=7, 7). H. Number of freezing bouts for GFP and CHR2 expressing mice during 13Hz laser stimulation in the test session. Number of freezing bouts for CHR2 mice predicted by Markov chain model is shown when using P_Fz/Fz_ fitted to laser stimulation data but keeping PAct/Act at the value prior to laser stimulation. This shows that the change in P_Fz/Fz_ is sufficient to predict the increase in freezing bouts. (Wilcoxon ranksum test: value=36.5: p=0.037; n=7,7).

These observations show that the 4Hz oscillation is more closely linked to termination compared to initiation of freezing bouts, suggesting a differential regulation of the two phenomena. In order to account for the difference between the two transitions, we turned to a probabilistic model of freezing behavior using a two-state Markov chain (freezing / active) which clearly separates freezing initiation and termination (Fig2F).

According to this model, at each time step, the animal is either in the freezing (Fz) or active (Act) state and randomly transitions to one of the two states according to probabilities P_Fz/Fz_ and P_Act/Act_ where P_Fz/Fz_ is the probability that freezing behavior will be maintained whereas P_Act/Act_ is the probability of staying in the active state (FigS4A). Both probabilities can be analytically derived from the average duration of freezing and active bouts respectively. Using the two probabilities obtained for each mouse, we simulated freezing time courses that allowed to almost perfectly predict the actual total time spent freezing found in the data both for GFP and CHR2 mice (FigS4B,C), This validates that the model captured the freezing-related behavior of the animals.

We then analyzed the effect of our optogenetic perturbation of the respiratory feedback within this framework. We found that only freezing bout duration was affected by our stimulation (Fig2G), which in this formalism is directly related to a single parameter P_Fz/Fz_. Therefore, perturbation of the OB specifically reduces P_Fz/Fz_ that controls freezing maintenance. On the contrary P_Act/Act_ that controls initiation, derived from active bout duration, is unaffected (Fig2G). These differences are also clear in the distributions of freezing and active bout durations (FigS4D).

By using this model, we can also predict the number of freezing bouts. Intuitively, one might imagine that modification of freezing could be due to either the modification of the duration or the number of freezing bouts. On the contrary, our model predicts that a selective drop in the probability of freezing maintenance (P_fz/Fz_) which makes freezing bouts shorter will lead to more opportunities to initiate freezing. Therefore, in spite of a constant probability to initiate freezing (1-P_act/Act_), the total number of freezing bouts should increase. This is exactly what is observed in our data and the value for each individual mouse can be predicted by the model when modifying only P_fz/Fz_ whilst holding P_act/Act_ constant (Fig2H). This further validates the model and supports the existence of two separate brain mechanisms controlling initiation and maintenance of freezing behaviour.

This analysis of optogenetic-induced changes of behavior therefore shows that only the maintenance of the freezing state is modified by OB perturbation. In the probabilistic framework of our model, termination and maintenance of freezing bouts are both controlled by the same parameter P_Fz/Fz_: the probability of maintaining freezing is the probability of not terminating it. Therefore, the specific modification of this parameter by OB perturbation is entirely consistent with the tighter locking of changes in OB 4Hz power to freezing offset compared to onset as described above.

After freezing initiation, the OB provides a feedback pathway for breathing related oscillations to impact downstream brain regions. Among the different areas that could link the bodily feedback with a modification of freezing behavior, the prefrontal cortex (PFC) is a good candidate structure since it plays a key role in emotional regulation and expression(26,27). More importantly it has been shown to display a prominent 4Hz oscillation during freezing reminiscent of what we describe here(28,29). We recorded LFP and single unit activity from the prelimbic area of the PFC (Fig S5) and found that during freezing the previously described 4Hz oscillation was highly coherent with OB activity (Fig3A,B,F,G,I)(23). Moreover granger causality shows that the OB clearly drives the PFC (Fig3C) in the 4Hz band and bulbectomy abolishes PFC 4Hz oscillations (Fig3D,H). This oscillation organizes neuronal activity, as shown by the phase-specific firing of around 45% of single units recorded to OB 4Hz oscillation (Fig3E). PFC single units display changes in firing during freezing (FigS6D) and we found that the individual activity of units was more strongly predictive of freezing vs active state at the freezing offset than onset (Fig3J,K). Strikingly, despite the profound changes in PFC activity dynamics induced by bulbectomy, no difference in overall firing rates was observed (FigS6A), suggesting that temporal organization of spiking activity rather that simple modification of rate coding is responsible for our observations. Our data therefore shows that feedback of respiratory-related 4Hz oscillations via the OB therefore organizes neural firing in the PFC and may allow this neural activity to regulate freezing maintenance.

**Figure 3.**
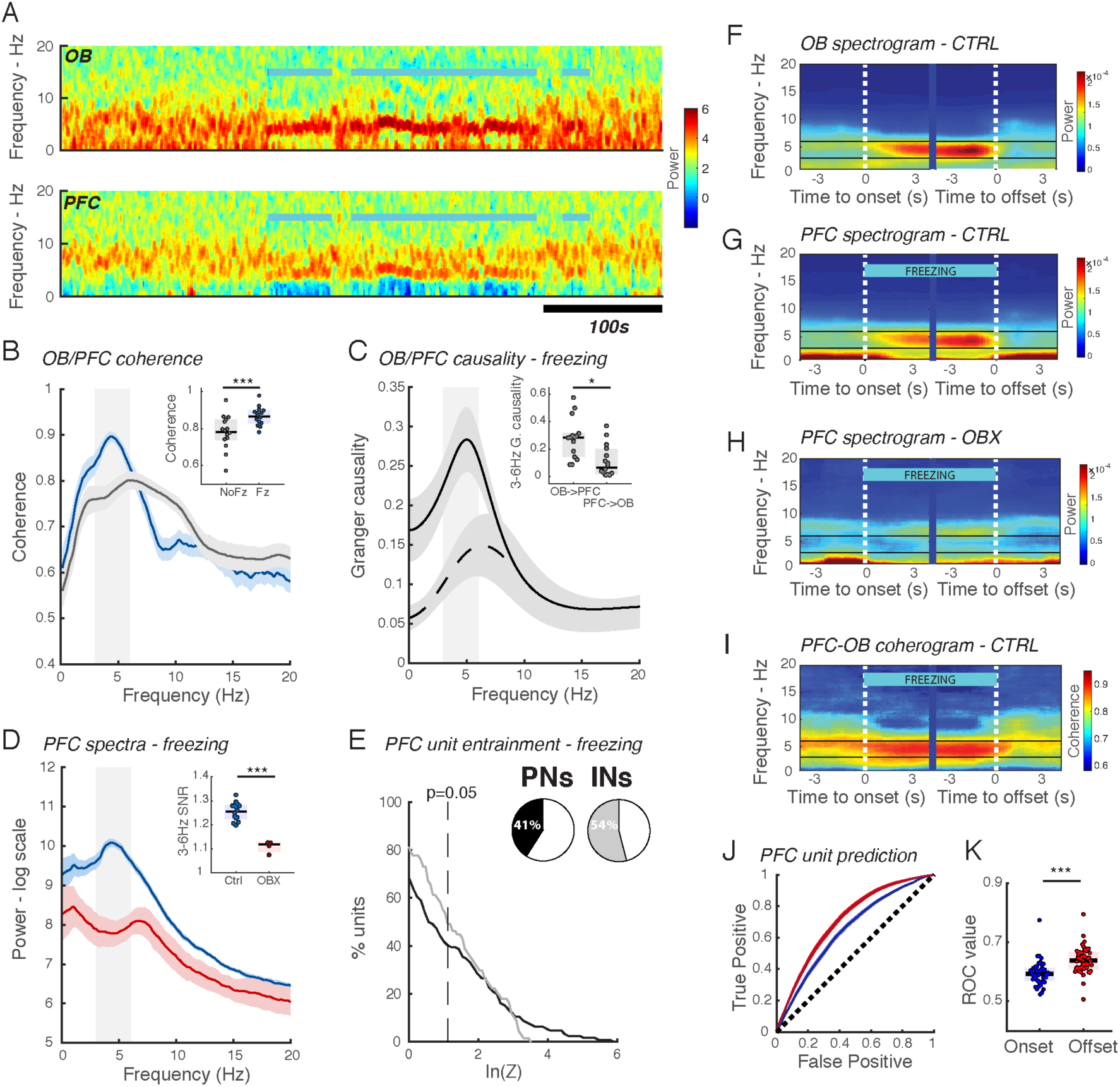
4Hz respiratory rhythm organizes neural firing in the prefrontal cortex during freezing. A. Representative evolution of spectral power in the olfactory bulb and the prefrontal cortex during a test session. Blue lines indicate the freezing periods. B. Averaged coherence between the olfactory bulb and prefrontal cortex during freezing (blue) and non-freezing (gray) periods. Error bars are SEM. Inset: average coherence of 3-6Hz band during freezing and non-freezing periods. (Paired Wilcoxon signed rank test: Signed Rank statistic=1, p=6.1e-5, n=15). C. Averaged granger causality during freezing from OB to PFC (full line) and from PFC to OB (dotted line). Error bars are sem. Inset: mean granger causality in the 3-6Hz band. (Paired Wilcoxon signed rank test: Signed Rank statistic=97, p=0.0353, n=15) D. Averaged power spectra during freezing in sham (blue) and bulbectomized (red) mice. Error bars are sem. Inset: signal-to-noise ratio of 3-6Hz band during freezing periods. (Wilcoxon rank sum test: Signed Rank statistic=143, p=8.4e-4, n=15 & n=4) E. Cumulative distribution of log-transformed Rayleigh’s test Z of PFC PNs and INs modulation by OB LFP. Inset shows percentage of significantly modulated PNs and INs using Rayleigh’s test with p=0.05. F. Averaged OB spectrogram triggered on freezing onset and offset in control mice. Spectrograms for each mouse are normalized to total power so that each mouse contributes equally to the average. Note the clear appearance of strong activity in the 4Hz band. G.H. As in D for the PFC in control (G) and bulbectomized (H) mice. Note the clear appearance of the 4Hz band in control mice whereas in bulbectomized mice there is no change of activity in this frequency band. I. Averaged PFC-OB coherogram triggered on freezing onset and offset in control mice. J. Receiver operating curves using firing of PFC single units to predict freezing or non-freezing state either at the onset or offset of freezing. Only units with a ROC value larger than 0.6 were included. The black line indicates the curve expected by chance. Error bars are sem. K. Area under the ROC curve freezing onset and offset. (Wilcoxon signed rank test: zval=4.83, p=1.35e-6, n=48 single units)

The respiratory rhythm can vary across multiple behavior states (19,30) over a large range of frequencies (2-12 Hz), raising the question of whether the impact on PFC activity during freezing is frequency-specific. We found that single units in the PFC were entrained more strongly and in larger numbers by the respiratory rhythm during freezing than during the active state (Fig4A, FigS7D,E see FigS7A-C for detailed methodology). Importantly, this result cannot be attributed to modification in spike number since any change in firing rate is uncorrelated with changes in phase modulation (Fig S6E). However, during freezing, changes in breathing frequency, depth and variability and many other factors such as neuromodulation could account for this change in phase locking.

**Figure 4.**
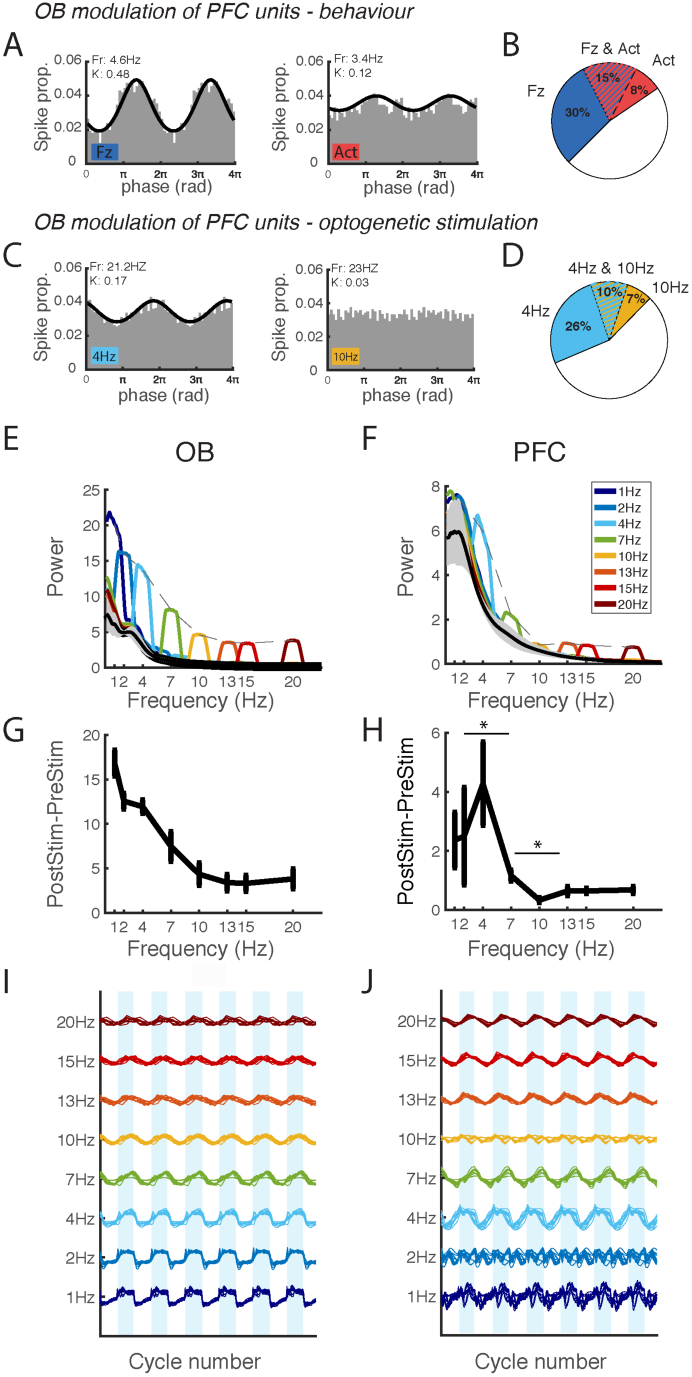
Optogenetic demonstration of frequency-specific tuning of the OB-PFC pathway to 4HZ. A.Phase histograms of an example PFC unit modulation by OB LFP showing clear gain in phase locking during freezing (left) relative to active (right) epochs. The firing rate and the relevant concentration coefficient of the neuron during the period is indicated on the graph. B. Pie chart showing the percentages of PFC units modulated by the OB LFP during active behaviour and during freezing using Rayleigh’s test with p=0.05. Chi2 test shows that the proportion of OB modulated neurons is increased for freezing state. (45% vs 23%, chi2stat=9.82, p=0.0017, n=100 units) C. Phase histograms of an example PFC unit’s modulation by OB LFP stimulation showing clear gain in phase locking during 4Hz (left) stimulation relative to 10Hz (right). D. Pie chart showing the percentages of PFC units modulated by the OB LFP during 4Hz and 10Hz stimulation using Rayleigh’s test with p=0.05. Chi2 test shows that the proportion of OB modulated neurons is increased for 4Hz stimulation. (chi2stat=18.82, p=1.43e-5, n=100 units). E.F. Average spectra in OB (E), PFC(F) before ChR2 stimulation in the OB (gray) and during stimulation at each frequency, color coded from lowest (blue) to highest (dark red). The dotted gray line shows the profile of frequency responses to stimulation in each structure. To ease comparison between mice with varying implantation depths, the spectra from each animal are normalized to total power. (n=4 mice) G. H Average difference in power between the pre-stimulation and stimulation period at each stimulation frequency for OB(G) and PFC(H).^∗^:4Hz and 10Hz power is respectively higher and lower than both neighboring frequencies in the PFC. (Friedman test, p=0.045 in all cases, n=4, error bars : sem) I.J LFP triggered on laser stimulation at each frequency showing a constant number of cycles for OB (I) and PFC (J). For each mouse, all responses are concatenated and z-scored to compare the relative amplitude at each frequency across individuals. (7 recordings sessions from 4 mice).

To parse the factors contributing to this change, we asked whether the PFC was equally receptive to all frequencies generated in the OB by optogenetically probing the circuit. We imposed a range of frequencies in the OB and measured their impact on the PFC activity. Stimulation was performed during sleep, a homogeneous behavioral state to avoid possible confounds due to modification of behaviour. OB LFP is successfully entrained at all frequencies by rhythmical optogenetic stimulation of GAD positive interneurons and shows a steady decrease in power as stimulation frequency increases with an 1/f profile (Fig4E,G,I). In the PFC, both the spectral power (Fig3F,H) and the amplitude of triggered LFP signals (Fig3J) show a peak response at 4Hz suggesting a resonance phenomenon and a total lack of response at 10Hz suggesting an antiresonance process.

This network property generalizes across brain states since it is observed both during NREM and REM sleep (FigS7F). Importantly these results are not a generic feature of neural responses to OB output because neither the OB nor the PiCx (the primary output structure of the OB) LFPs show this response profile (FigS8B,C,D). Instead, the PiCx displays a clear resonance at 20Hz. The entrainment of single units recorded in the PFC (n=169) showed the same pattern of preferential entrainment to 4Hz compared to 10Hz (Fig4C,D, FigS7G). Changes in PFC units firing rates and therefore excitability cannot explain this change (FigS6D,E). These results demonstrate that the OB-PFC dialog is specifically tuned to amplify communication at 4Hz, the frequency associated with freezing, and to impede communication at 10Hz, the frequency associated with active exploration.

To summarize, we show that freezing is associated with regular 4Hz breathing in mice, this oscillation feeds back to the brain via the OB and both the behavioral effects of optogenetic perturbation and analysis of electrophysiological data argue that it then specifically regulates the maintenance of freezing behavior. The most likely pathway of action is through the PFC which is optimally entrained by the specific frequency of 4Hz breathing associated with fear behavior and in which neuronal firing patterns are consistent with a role in regulating freezing maintenance. The role of the PFC is therefore to determine whether the current coping strategy will be continued or interrupted which can reconcile its role in emotional regulation with its role in relation to the temporal organization of behavior and perseverance in rule switching tasks(31,32).

Thus, we show that bodily feedback plays both an active and direct role in emotional states. First, it is active in that it specifically regulates the duration of emotional states. Second, it is direct in that its influence is not mediated by the representation of interoceptive signals but should instead be viewed as part of the computation itself since the breathing rhythm generates dynamics that organize neural activity. This provides some of the strongest support to date for theories of radical embodied cognition(33–35).

Taken together, our findings thus lead us to propose a causal chain of emotional generation that involves two major steps. The process begins with the initiation of changes in behavior and physiology by the classical pathways of the amygdala and periaqueductal gray. Indeed, stimulation of these areas can robustly initiate freezing behavior(36–38) and amygdala ablation modifies bout initiation(39). Moreover, these structures are well known to modify breathing for example via the nucleus retroambiguus (40,41). Once initiated, the regular breathing will then entrain the OB, organize neuronal activity in the PFC and modulate the probability of freezing maintenance (FigS1).

This model enables the James-Lange theory to be reconciled with the objections raised by Walter Cannon and subsequent reports of inconsistent changes in emotion under impaired bodily feedback(9–12). Indeed, in our framework the bodily expression of emotions is part of a regulatory feedback loop which is crucial for the stabilization of the emotional state, in spite of an independent process of initiation. We therefore propose that the crucial factor determining whether an emotive situation gives rise to a fully-fledged emotion is sufficient maintenance of this state via this feedback loop but that, in pathological cases, it could emerge from repeated initiation events. This leads to the clear prediction that careful investigation of human emotion in cases of privation from bodily feedback, such as in cases of spinal cord injury, should reveal that remnant emotional experiences will be of shorter duration. The umbrella term ‘emotion’ must therefore be parsed into three sequential facets: a sub-cortical emotion leads to a bodily emotion which must feedback to the cortex for this state to last long enough to become a full cognitive – and perhaps conscious – emotion.

## Acknowledgments

We thank L. Roux and A. Sirota. for discussions, and A. Peyrache for discussion and critical readings of an earlier version of the manuscript.

## Author contributions

S.B. and J.M.L. performed the experiments and analysed the data. M.M.L. and G.d.L. did several experiments included in the dataset. C.B. helped for the behavioural experiments. H.G. performed the histology. S.B. and K.B. wrote the manuscript with the help of J.M.L. and C.H..

## Data and materials availability

All data and code can be made available on request to the corresponding author.

## Materials and methods

### Subjects

Mice were housed in an animal facility (08:00–20:00 light), one per cage after surgery. All behavioural experiments were performed in accordance with the official European guidelines for the care and use of laboratory animals (86/609/EEC) and in accordance with the Policies of the French Committee of Ethics (Decrees n° 87–848 and n° 2001–464). Animal housing facility of the laboratory where experiments were made is fully accredited by the French Direction of Veterinary Services (B-75-05-24, 18 May 2010). Animal surgeries and experimentations were authorized by the French Direction of Veterinary Services for K.B. (14-43).

### Electrode implantation and virus injection

C57Bl6 male mice between 3 and 6 months old were implanted under deep anesthesia with a xylazine (10m g/kg)-ketamine(100 mg/kg) mixture. Electrodes were placed in the right olfactory bulb (AP +4, ML +0.5, DV –1.5), the right CA1 hippocampal layer (AP −2.2, ML +2.0, DV –1.0) and the right prefrontal cortex (AP +2.1, ML +0.5, DV –0.5).

A total of 17 GAD-Cre-line mice were injected with AAV5-ChR2 virus (10 mice) and control AAV5-GFP virus (7 mice) in bilateral olfactory bulbs. A range of injection sites were used to obtain optimal to the glomerular layer (GL) since GL interneurons are thought to generate the OB slow oscillations whereas to the granular layer interneurons are involved in OB fast oscillations(1). Injection sites are listed below, we found no differences in electrophysiological response so all mice were pooled. Virus injections were performed with a glass capillary at a rate of 150 nL/min and followed by a 10min pause. 3 weeks after injection, mice were bilaterally implanted with home-made optic fibres(2).

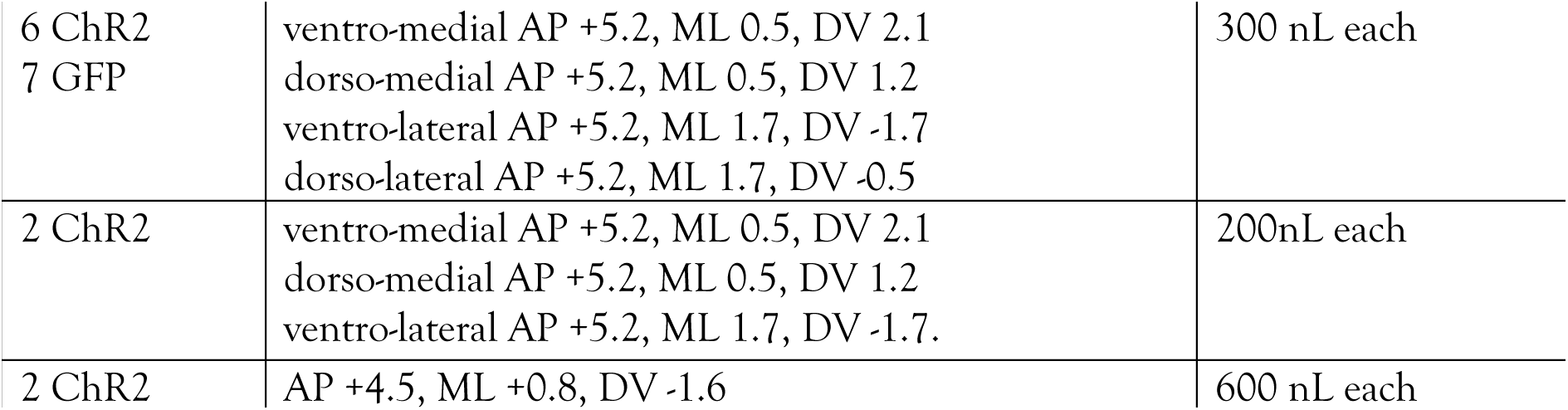

During recovery from surgery (minimum 1 week) and during all experiments, mice received food and water ad libitum.

### Bulbectomy

Animals were randomly assigned to the bulbectomy and sham-operation groups. The animal was placed in a stereotaxic frame and two burr holes were drilled on either side of the midline of the frontal bone above the olfactory bulbs. The bulbs were aspirated with a micro-pipette. The cavity was packed with bonewax, the holes were covered with dental cement and the skin sutured. Sham-operated mice were treated similarly, except that the holes were not drilled. After the surgery, animals were housed one per cage. Animals were handled and weighed daily during a 7-day recovery period.

### Electrophysiological recordings & spike sorting

Signals from all electrodes were recorded using an Intan Technologies amplifier chip (RHD2216, sampling rate 20 KHz). Local field potentials were sampled and stored at 1,250 Hz. Analyses were performed with custom made Matlab programs, based on generic code that can be downloaded at www.battaglia.nl/computing/ and http://fmatoolbox.sourceforge.net/.

Local field potentials were recorded using tungsten wires with PFA isolation (0.002″ bare) and single units using home-made tetrodes formed by twisting insulated Nichrome Wire (0.001” bare). Tetrodes were plated to reduce impedances to around 100kΩ using gold solution (Neuralynx).

For spike sorting, extracted waveforms were sorted via a semi-automated cluster cutting procedure using KlustaKwik and Klusters (http://neurosuite.sourceforge.net/) Recordings were visualized and processed using NeuroScope and NDManager (http://neurosuite.sourceforge.net/).

### Fear conditioning

Habituation to sounds and fear conditioning took place in context A consisting of a cylindrical transparent Plexiglas box in a black environment with a shock grid floor and cleaned with ethanol (70%) before and after each session. Habituation to test environment and the test session were performed in context B consisting of square transparent Plexiglas walls with a grey plastic floor placed in a white environment and cleaned with acetic acid (1%) before and after each session. On day 1, mice were habituated to the test environment without sounds. On day 2, mice were submitted to a habituation session during which they the CS- and of the CS+ were presented 4 times each (total CS duration, 30 s; consisting of 50-ms pips at 0.9 Hz repeated 27 times, 2 ms rise and fall; pip frequency, 7.5 kHz or white-noise, 80 dB sound pressure level). Discriminative fear conditioning was performed on the same day by pairing the CS+ with a US (1-s foot-shock, 0.6 mA, 8 CS+ US pairings; inter-trial intervals, 20–180 s). The onset of the US coincided with the offset of the CS+. The CS-was presented after each CS+ US association but was never reinforced (8 CS-presentations; inter-trial intervals, 20–180 s). On day 3, conditioned mice were submitted to a test session during which they received 4 and 12 presentations of the CS- and CS+, respectively.

To score freezing behaviour animals, were tracked using a home-made automatic tracking system that calculated the instantaneous position of the animal and the quantity of movement defined as the pixel-wise difference between two consecutive frames. The animals were considered to be freezing if the quantity of movement was below a manually-set threshold for at least 2s. Freezing behaviour was quantified during blocks of CS as stated in each figure by calculating the percentage of time spent freezing during the sound and the subsequent 30s. Some animals were equipped with an Intan RHD2132 board which includes as 3-axis accelerometer which gave access to the animal’s acceleration with high temporal resolution.

### Plethysmography

Respiration was recorded with a whole-body plethysmograph (Emka Technologies) with continuous air flow. The pressure signal was amplified via an omnetics headstage and recorded using the same Intan system as the electrophysiological signals to insure synchrony. Breathing frequency is defined as the inverse of the time between two inspirations and the tidal volume is the integral of the negative signal. The system was calibrated to convert voltage fluctuations into mL/s by manually injecting known volumes of air at different speeds.

### Optogenetic stimulation

#### Fear conditioning experiment

On day 1, during the habituation session, mice received 13Hz blue light stimulations (45s). The stimulation intensity was chosen to see clear LFP oscillations in the OB and the PiCx, an output structure. No stimulation was delivered on day 2, during the habitation to sounds and the conditioning phase. During the test session on day 3, mice first received 3 shorts bouts of light stimulation (10s, inter-stim interval 10s) before the onset of the first CS. No stimulation was delivered during the CS- and the first pair of CS+. 13Hz blue light stimulations were delivered during subsequent CS+ (45s, locked at the CS+ onset, i.e. extending 15s after the end of each sound).

#### Sleep stimulation experiment

After an initial light-phase day of baseline recording in their homecage to habituate to sleeping with electrophysiological and optic cables, mice were tracked using homemade Matlab software to identify periods of prolonged immobility corresponding to sleep. During these periods, periodic stimulation of ChR2 was delivered at a randomly selected frequency take from 1,2,4,7,10,13,15 or 20Hz. Stimulations were separated by at least 5min. Light intensity was calibrated to be at 15mW/mm2.

All experiments used 473nm light, delivered using a Doric Lense rotary joint and patch cords to a fiber of 0.37 numeric aperture.

### Histological analysis

After completion of the experiments, mice were deeply anesthetized with ketamine/xylazine solution (10% /1%). With the electrodes left *in situ*, the animals were perfused transcardially with 50 ml saline, followed by 50 ml of PFA (4 g/100 mL). Brains were extracted and placed in PFA for postfixation for 24 h, transferred to PBS for at least 48 h, and then cut into 50-μm-thick sections using a freezing microtome. Slides were mounted and stained with hard set vectashield mounting medium with DAPI (Vectorlabs).

### Spectral and coherence analysis

Electrophysiological data were analysed with custom-written Matlab (The MathWorks Inc., Natick, MA, 2000) codes that made use of the Chronux toolbox (http://www.chronux.org). Spectrograms and coherograms were calculated with multi-taper Fourier analysis to diminish finite windowing effects. For all analysis, we applied 5 tapers and a window size of 3s with a 0.1s shift between bins. Changes in power of specific bands were calculated only after identification of a peak in the power spectrum (3-6Hz for the respiratory rhythm) in at least one condition using the signal to noise ratio defined as the ratio between power in the band of interest to the power in the rest of the spectrum.

To quantify the relationship between behaviour (freezing or not freezing) and power in the 3-6Hz band, we used Receiver Operator Characteristic curves(3). This analysis quantifies the ability of an ideal observer to predict whether an animal is freezing or not based purely on the instantaneous power in the 3-6Hz band. The ROC analysis presents the advantage over other evaluations of classifier accuracy of being insensitive to class distribution, that is the respective number of freezing and non-freezing bins in our case which can often be strongly skewed depending on the behavior of the animal. However, we restricted our analysis to animals presenting at least 10s of freezing and 10s of non-freezing behavior (n=12/15). When analyzing specifically onset and offset prediction, mice were further restricted for having a sufficient number of freezing bouts longer than 6s (n=11/15).

In our case, the observed discriminates the two behavioral states by placing a threshold (z) on 2s-binned spectral power (r) in the 3-6Hz band below which the corresponding bin is classified as ‘freezing’ and above which it is classified as ‘non-freezing’. The performance of this procedure can be fully determined by two parameters:

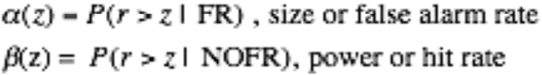

Plotting α and β for increasing values of z yields the ROC curve and the area under this curve curve (ROC value) represents the probability that an ideal observer can discriminate between a 2s period of freezing or no freezing based on the spectral power recoded at that time: it is equal to 0.5 if the spectral power carries no information about the behaviour and equals 1 it is perfectly predictive. We took 0.6 as a threshold for minimal prediction value, as suggested in (4).

### Phase analysis of LFP signals

Instantaneous phase of the OB LFP during freezing was derived by applying the Hilbert transform to the filtered signal. To estimate the instantaneous phase of signals throughout the behavioural session with highly variable frequencies, particularly for the OB respiratory rhythm that varies from 2 to 12Hz, we applied a method derived from(5) as filtering in frequency bands to estimate instantaneous phase would fail to capture these dynamic changes. The LFP signal was filtered between 1Hz and 20Hz and local peaks and troughs were detected that exceeded 0.5 times the standard deviation of the signal. After this first pass that produced a list of times of maxima, successive troughs or peaks were detected. If a peak or trough was identified between the two extrema by applying a softer threshold (0.2 times the std), this point was added, otherwise the smaller of the two maxima was deleted. This second pass guaranteed the succession of troughs and peaks in the time series. The parameters used were empirically established after extensive visualization of the extrema detection on both HPC and OB signals. The phase of the signal was then defined by interpolating between the peak (0°) and the trough (180°). To evaluate the quality of the method for capturing the instantaneous variations in the signal’s frequency, a test signal was reconstructed using the instantaneous phase and the amplitude of the detected extrema. The mean square error was calculated to estimate the quality of the reconstruction (Fig S7C).

### Phase modulation analysis of units

Only test sessions with more than 10s of freezing were used in this analysis. Using the methods described in the previous section, each spike of each unit was attributed its corresponding phase and the distribution of these phases was tested for uniformity. Given the assymetric shape of OB oscillations, the phase distributions were corrected as described in(6). To test the significance of phase modulation we used the standard Rayleigh test for uniformity with a threshold of p=0.05. The relevant statistic used for the test is the variance-stabilized log(Z)=R^2/n where R is the resultant length and n the number of spikes.

When comparing the modulation of units by OB LFP in active and freezing states (Fig 4A,B), we corrected for period duration by only analyzing the period of highest activity of corresponding length to the freezing period. We also corrected for changes in firing rate by subsampling spikes using the bootstrapping method in the period with the highest firing rate. We compared the strength of modulation using two parameters: the frequently used concentration coefficient kappa and the pairwise phase consistency(7) which is an unbiased measure (Fig S7DE). To estimate the phase preference of PFC units to the respiratory rhythm using OB LFPs, we corrected the phase of modulated units by the average phase difference between PFC and OB recordings. This allowed us to avoid confounds due to phase changes with depth in the OB.

### Cell type classification

For each unit, we used three parameters taken from the literature(8,9) to cluster waveforms into two clusters : duration at half-amplitude, trough to peak latency and the asymmetry index (positive ratio of the difference between right and left baseline-to-peak amplitudes and their sum) (Fig S5C,D). To cluster the units used in the study we grouped them with a larger set of units (629) recorded from other studies in the laboratory to insure more robust clustering. All together these units were clustered into 74% putative pyramidal neurons and 26%. Putative pyramidal neurons were broader that putative interneurons and as expected had lower average firing rate (5.5Hz +/−0.9 s.e.m vs 14.7Hz +/−1.9 s.e.m).

### Markov model of behavior

To apply the two-step Markov chain model, the continuous time data is chunked into steps of 2s length. This time bin was chosen to optimize the model’s predictions of total freezing time and frequency of bout initiations. Bout duration, expressed in number of time bins, is related to the probability to stay in that state by the relation:

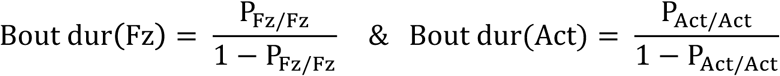

These relations imply that both parameters of the model can be directly derived from the measured mean bout duration of freezing and active states. To validate the model, we then simulated the model 100 times to estimate two variables that could be derived from the data: total freezing duration and frequency of freezing bout initiations. Unlike average bout duration, these variables depend on both parameters of the model and therefore do not allow to disambiguate which of the two processes is affected by a given manipulation.

### Modulation index

To evaluate changes in a given parameter X (firing rate, concentration coefficient) at the level of the individual unit, we define the modulation index to compare situation 1 and 2 for each neuron as:

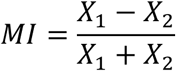

### Statistics

For each statistical analysis provided in the manuscript, the Kolmogorov–Smirnov normality test was first performed on the data. If the data failed to meet the normality criterion, statistics relied on non-parametric tests. We therefore represent the median and quartiles of data in boxplots in all figures, in accordance with the use of non-parametric tests.

For comparison of proportions, the standard chi2 test was used.

Two-way mixed repeated anova was performed using GraphPad Prism, GraphPad Software, La Jolla California USA.

Ranksum and signed rank : we report the signed rank statistic if the number of replicates is too weak to provide the normal Z statistic

## Supplementary Materials

**Fig. S1.**
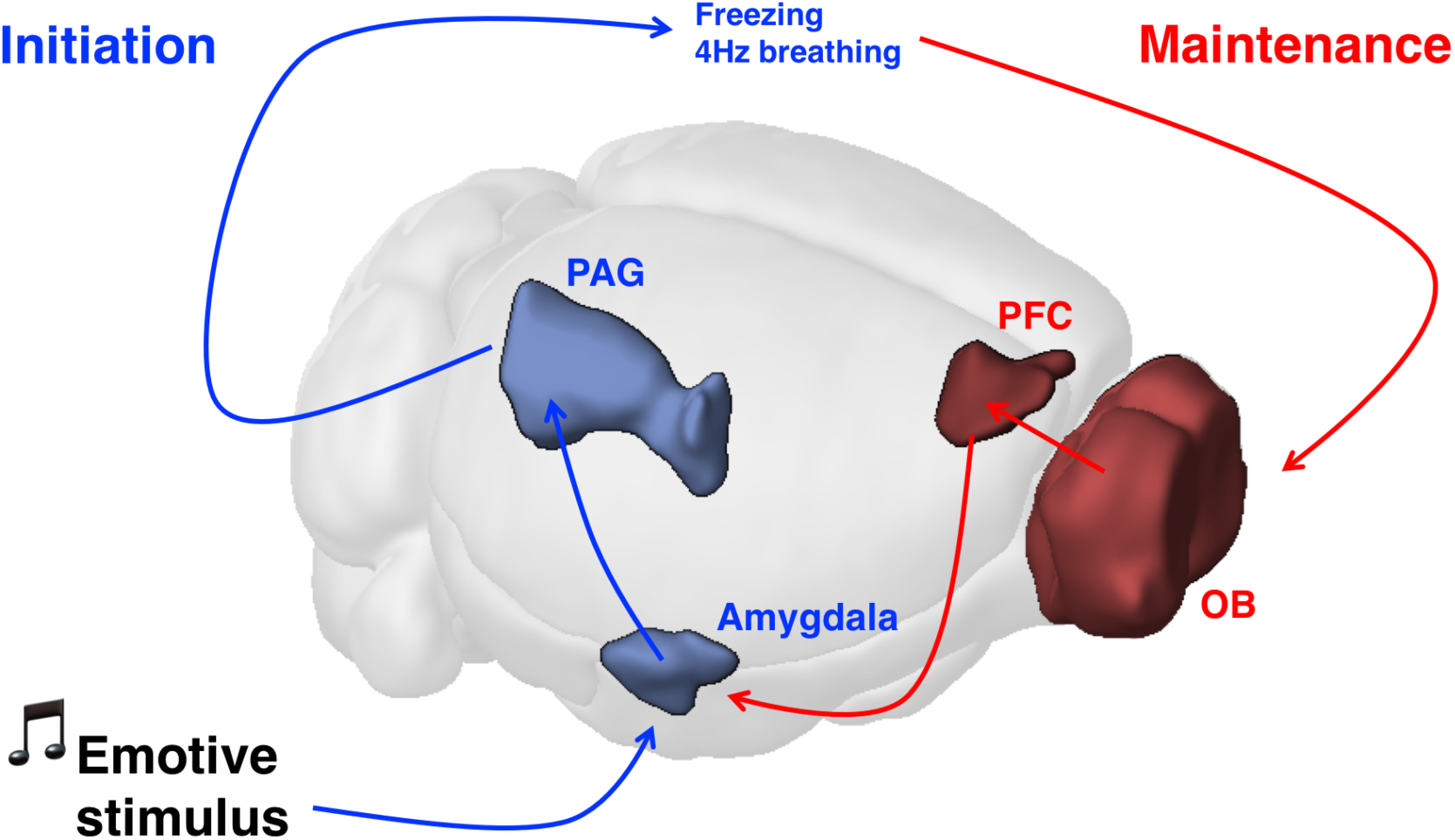
Model of brain-body-brain feedback loop of emotional initiation and maintenance. An emotive stimulus evokes responses in the amygdala and Periaqueductal Gray (PAG) which lead to changes in behavior (freezing) and somatic physiology (breathing). The 4Hz breathing entrains the OB which in turn organizes neural firing in the PFC which allows to maintain the ongoing behavior, perhaps via its connections with the amygdala for example.

**Fig. S2.**
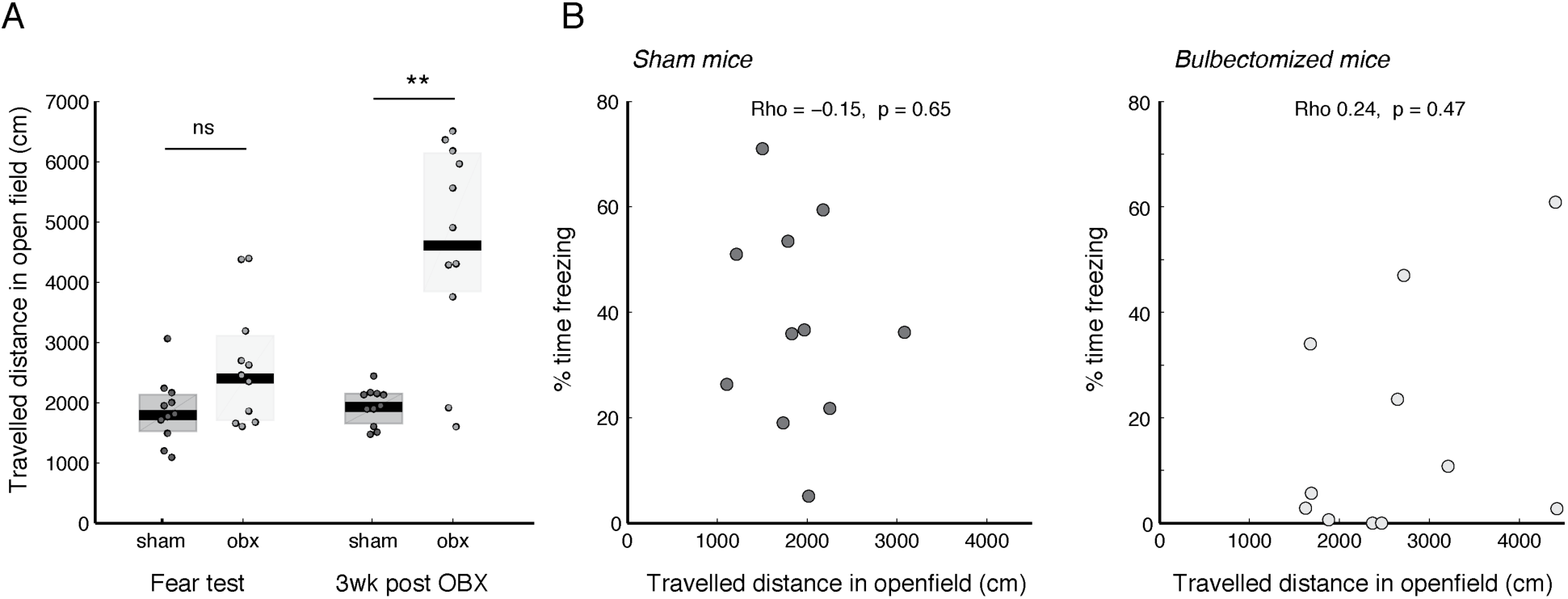
A. Travelled distance in the open field test after 10min for sham and bulbectomized (obx) mice, at two time points: the day following the fear test (Wilcoxon rank sum test test, zval=-1.773, p=0.076, n=11 & 11 mice) and one week later, i.e. 3 weeks after bulbectomy (Wilcoxon rank sum test, zval=-2.955, p=0.003, n=11 & 11 mice). B. No correlation between freezing level and the travelled distance in the open field test following the fear test, either for sham (left, Spearman correlation coefficient Rho=-0.15, p=0.65) or for bulbectomized mice (right, Rho = 0.24, p=0.47). This is consistent with the fact that removing the 2 bulbectomized mice with a high travelled distance in open field on the day post-fear does not modify the difference of freezing during CS+ (Two-way mixed anova: group (p=0.006, F=9.702) x CS block (p<0.0001, F=10.86), interaction (p=0.029, F=3.230). Post-hoc Wilcoxon ranksum on each individual block : (zval=1.4463, 2.2332, 2.6296, 2.2389; p=0.148, 0.025, 0.0075, 0.0174). Thus, decrease in freezing in bulbectomized mice cannot be explained by a change in locomotor activity since hyperactivity does not develop until three weeks later and the overall level of locomotion does not correlate with percent of time freezing.

**Fig. S3.**
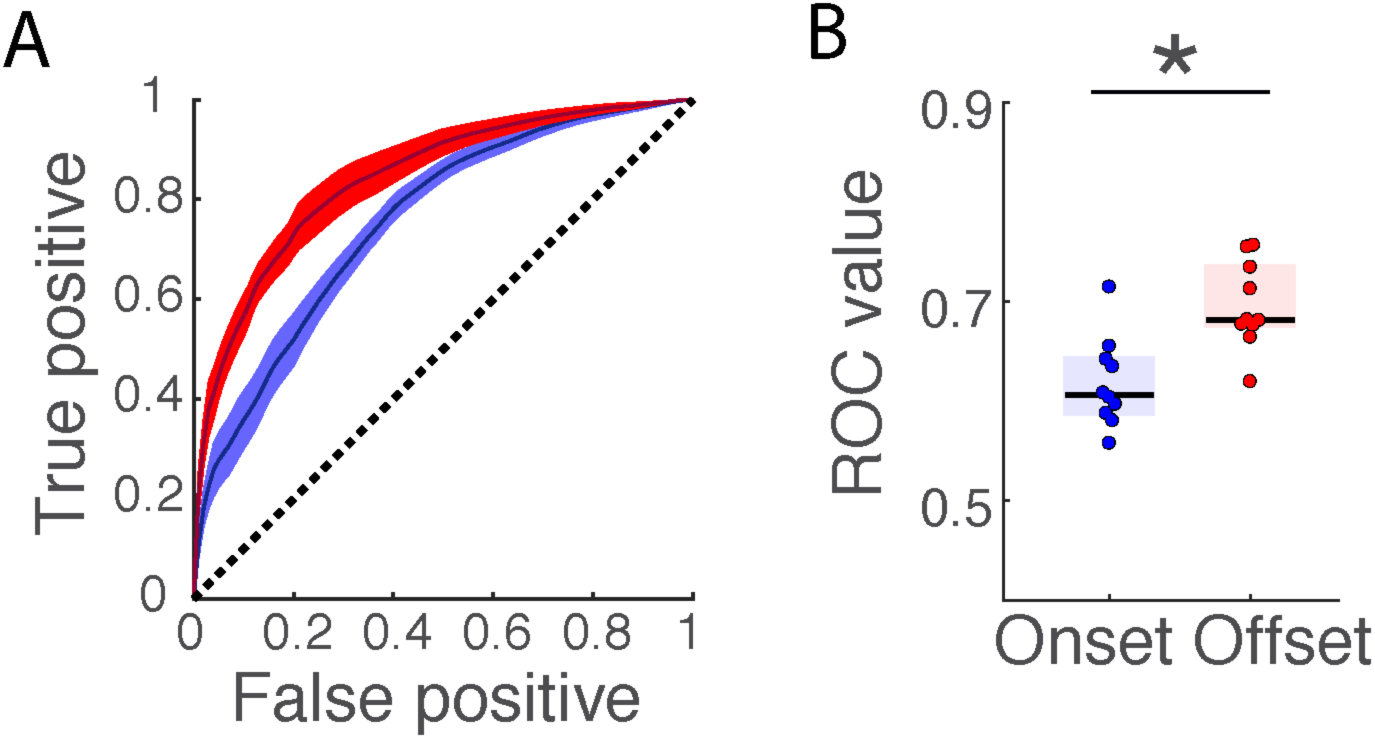
A. Receiver operating curves using power in the 4Hz band to predict freezing or non-freezing state either at the onset or offset of freezing. The black line indicates the curve expected by chance. Error bars are sem. B. Area under the ROC curve freezing onset and offset. (Wilcoxon signed rank test: Signed Rank statistic=55, p=0.0019, n=10 mice)

**Fig. S4.**
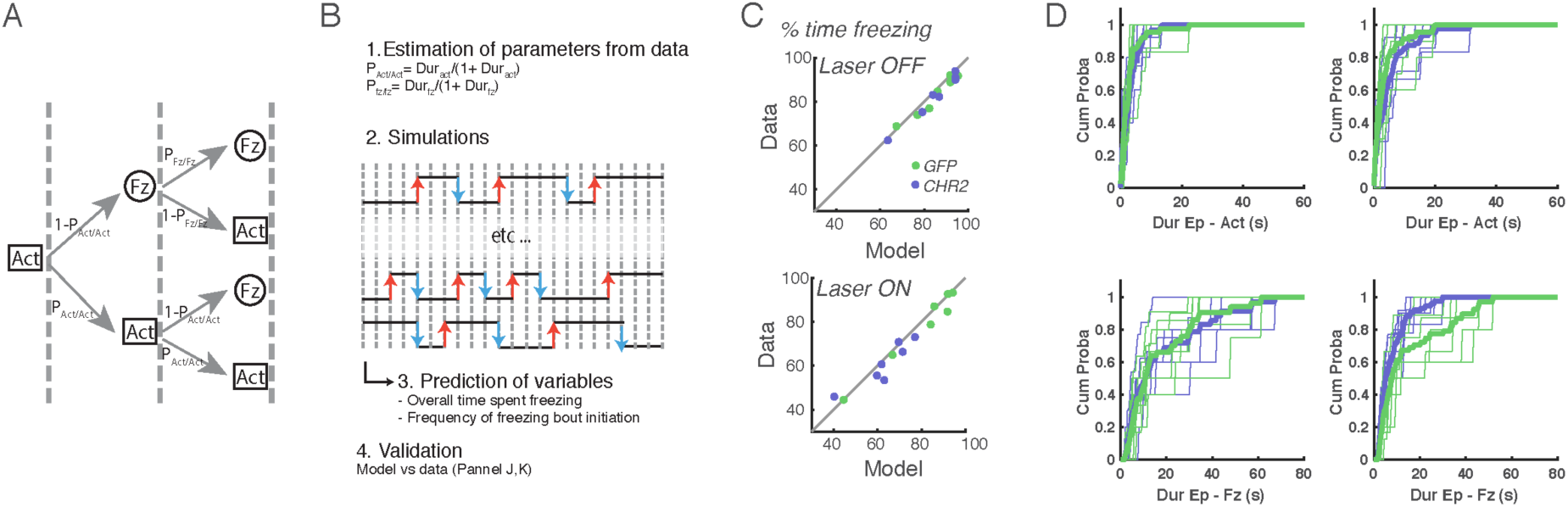
A. Scheme of time steps that follow the Markov model used to model freezing. At each time step, depending on the current state (Fz or Act), the next state is randomly selected according to fixed probabilities. B. Scheme of model derivation from data and testing. The two probabilities (P_Act/Act_ and P_Fz/Fz_) are directly estimated from the average active and freezing bout durations. To establish the quality of the model, two other parameters not used to calibrate the model, the percent of time spent freezing and the frequency of freezing bouts, are estimated from the stimulations to be compared to data. C. Correlation of data with simulation-based prediction of the Markov model for the total percent of time spent freezing. Note that both before and during laser stimulation all points lie close to the identity line shown in black, indicating an excellent prediction by the model. (Correlation coefficients: laser off: R=0.94, p=5.12e-7; laser on: R=0.99, p=7.5e-12; n=7 7 mice) D. Cumulative distribution of active and freezing bout lengths for GFP and CHR2 expressing mice during CS+ presentations before (left) and during (right) laser 13HZ stimulation. Note that without stimulation the distributions are nearly identical. During stimulation, although active bout duration curves remain similar, there is an upwards shift of the freezing bout duration curve for CHR2-expressing mice relative to their GFP controls, indicating a reduction in freezing bout duration.

**Fig. S5.**
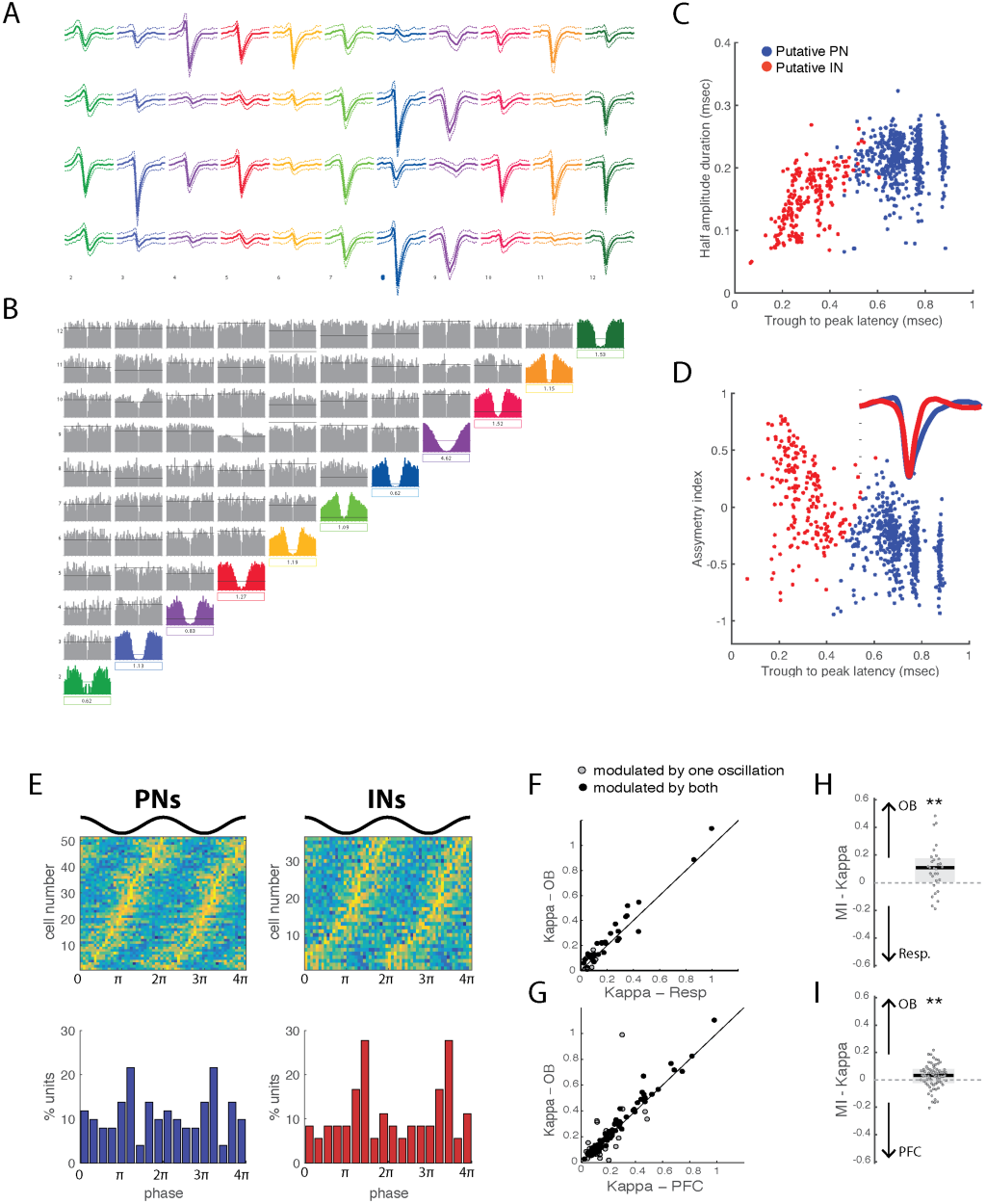
A. B. Example spike sorting sessions showing mean waveforms on each wire of the tetrode and cross and auto-correlograms for each identified single unit. C. D Waveforms were clustered using three waveform characteristics: duration at half-amplitude, trough to peak latency and the asymmetry index (positive ratio of the difference between right and left baseline-to-peak amplitudes and their sum) Putative pyramidal neurons (blue) and interneurons (red) were clustered using k-means algorithm. Inset shows mean waveforms of all pyramidal neurons (blue) and interneurons (red). E. Phase distribution of all significantly modulated PFC PNs and INs by OB LFP (top) and distribution of preferred phases for all significantly modulated PNs and INs (bottom). F. Correlation of the von Mises concentration coefficient (K) of PFC units during freezing relative to respiration and the 4Hz oscillation in the OB LFP. Significantly modulated units for either or both signals are shown. G. As for F but for 4Hz oscillation in the PFC vs the OB LFP. H. Modulation index of the von Mises concentration coefficient (K) showing significantly stronger modulation for the OB signal than the respiration signal for units that are modulated by both signals. (Wilcoxon rank sum test, p=0.0032, zval=2.95, n=30 units). I. As for H but for 4Hz oscillation in the PFC vs the OB LFP. (Wilcoxon rank sum test, p=0.0029, zval=2.98, n=76 units).

**Fig. S6.**
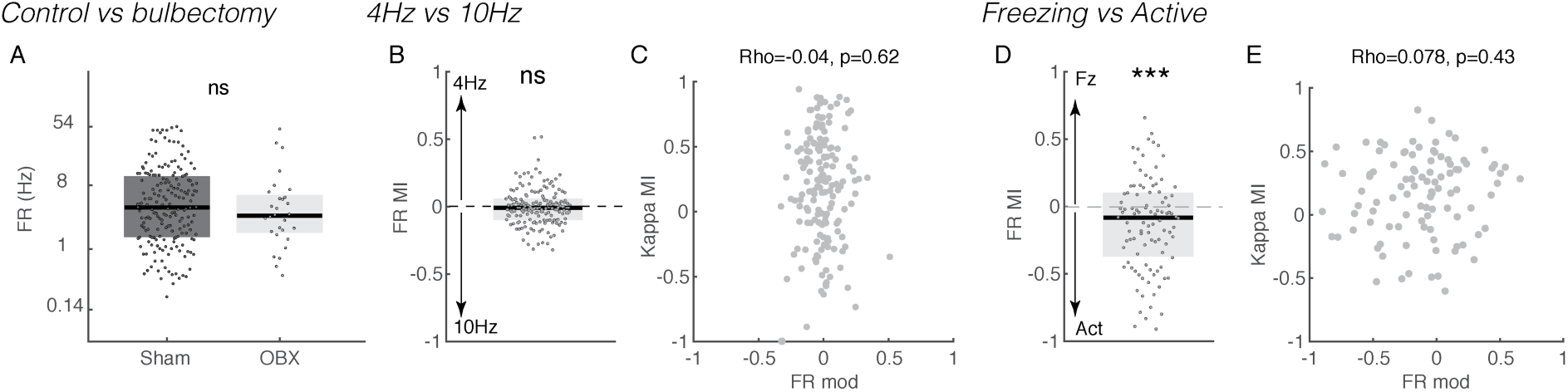
A. Firing rates of PFC units in control and bulbectomized mice, showing no difference in overall level of activity. (Wilcoxon rank sum test, zval=0.8986, p=0.3689, n=191 units in 13 control mice, 29 units in 3 bulbectomized mice). B. Modulation index of firing rate of all neurons between 4Hz stimulation periods and 10z stimulation periods, showing no overall change. (One-sample two-sided Wilcoxon signed rank test with 0: p=0.24, zval=-1.1813, n=167 units) C. Modulation index of concentration coefficient Kappa as a function of the modulation index of firing rate of all neurons showing the absence of correlation. (Spearman correlation coefficient: R=-0.04, p=0.62, n=167 units). D. Modulation index of firing rate of all neurons between freezing and active periods showing that on average units decrease their firing rate during freezing. (One-sample two-sided Wilcoxon signed rank test with 0: zval=-3.4314, p=6.0e-04, n=100 units) E. Modulation index of concentration coefficient Kappa as a function of the modulation index of firing rate of all neurons showing the absence of correlation. (Spearman correlation coefficient: R=0.078, p=0.43, n=100 units).

**Fig. S7.**
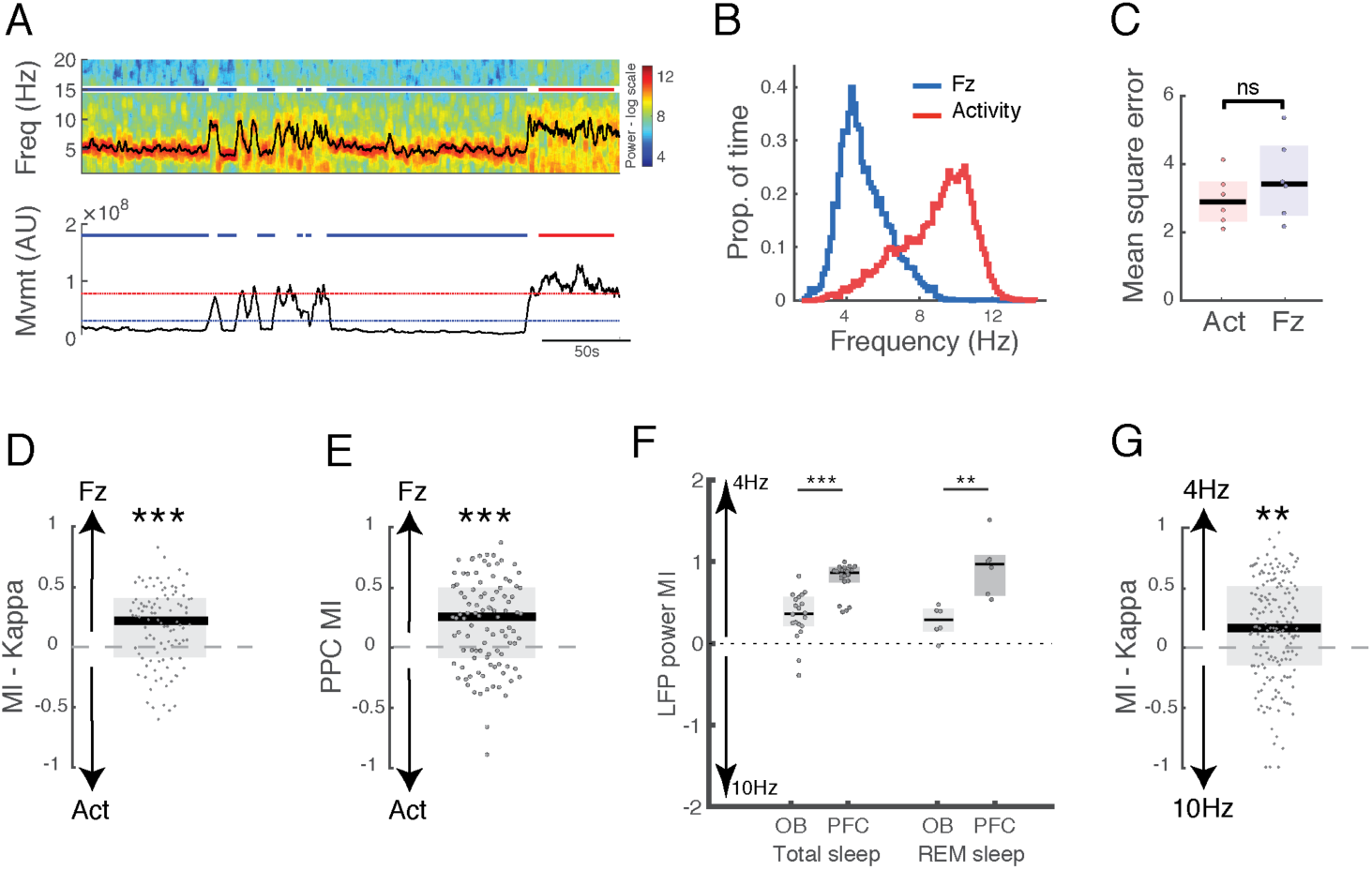
A. Example spectrogram of OB activity showing the instantaneous frequency estimated using the peak-to-peak phase estimation method (black line). Note that the instantaneous frequency closely tracks the strongest band of the spectrogram despite changes in frequency. Below the quantity of movement illustrates the definition of freezing (below the blue line) and active (above the red line) epochs. The active threshold was adjusted to have equivalent total duration of data for both epochs. Red and blue lines represent active and freezing epochs respectively. B. Distribution of instantaneous frequencies of OB LFP during freezing and active periods. Freezing frequencies peak around 4Hz whereas active frequencies show a broader distribution around 10Hz. Note that these frequencies are slightly higher the breathing frequencies shown in Fig1 because in the restricted volume of the pletysmograph mice could not attain high speeds and here only time points of active exploration are used, not merely all time points outside of freezing which may include quiet wakefulness. C. Mean square error between the LFP signal in the OB and the reconstructed signal using interpolated phase estimated using the peak-to-peak method. Error rates were not significantly different during the active and freezing periods, indicating that this method allows us to track the phase of the despite changing frequencies. (Paired Wilcoxon signed rank test: signed rank statistic: 5; p=0.31, n=6 mice). D. Modulation index of the concentration coefficient for PFC units relative to OB LFP showing a significant increase in spiking phase modulation during freezing relative to active epochs. Arrows on the left indicate how the figure can be interpreted: positive values indicate an increase in phase modulation during freezing behavior. (One-sample two-sided Wilcoxon signed rank test with mean 0: zval= 4.6349, p=3.5718e-06, n=100 units in 6 mice). E. Modulation index of the pairwise phase consistency index for PFC units relative to OB LFP showing a significant increase in spiking phase modulation during freezing relative to active epochs for the OB signal only. This measure is robust to changes in firing rate. Arrows on the left indicate how the figure can be interpreted: positive values indicate an increase in phase modulation during freezing behaviour. (One-sample two-sided Wilcoxon signed rank test with 0: zval= 5.06, p=4.2e-07, n=100 units) F. Modulation index of OB and PFC responses to optogenetic stimulation of the OB at 4Hz relative to 10Hz during total sleep (left) and REM sleep only (right). Note that there are fewer points in the REM sleep quantification because data was restricted to sessions with sufficient number of stimulations during this short sleep epoch. (Two-sample two-sided Wilcoxon ranksum test: zval=-4.96, p=1.1e-4, n=20; ranksum value=21, p=0.0022, n=6) G. Modulation index of the concentration coefficient Kappa for PFC units relative to laser stimulation, showing a significant increase in spiking phase modulation during 4Hz stimulation relative to 10Hz stimulation. Arrows on the left indicate how the figure can be interpreted: positive values indicate an increase in phase modulation during 4Hz. One-sample two-sided Wilcoxon signed rank test with mean 0: zval=5.4894, p=4e-8, n=100 units).

**Fig. S8.**
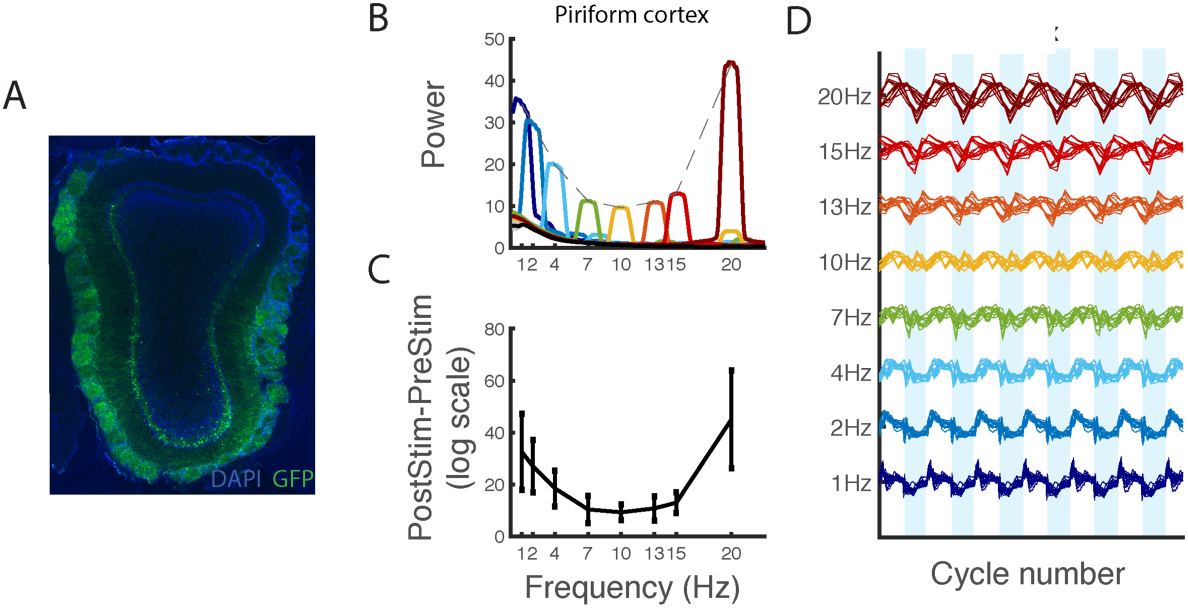
A. OB histology of ChR2 injected mice. Note the presence of ChR2+ cells in the glomerular layer. B. Average spectra in PiCx before ChR2 stimulation in the OB (gray) and during stimulation at each frequency, color coded from lowest (blue) to highest (dark red). The dotted gray line shows the profile of frequency responses to stimulation in each structure. To ease comparison between mice with varying implantation depths, the spectra from each animal are normalized to total power. (n=4 mice) C. Average difference in power between the pre-stimulation and stimulation period at each stimulation frequency in PiCx. (n=4 mice, error bars : sem) D LFP triggered on laser stimulation at each frequency showing a constant number of cycles in PiCx. For each mouse, all responses are concatenated and z-scored to compare the relative amplitude at each frequency across individuals. (7 recordings sessions from 4 mice).

